# Plasma donation induces a protein expression profile shift in circulating human blood

**DOI:** 10.1101/2021.10.30.466597

**Authors:** Rob Flickenger

## Abstract

Commercial plasma donation yields a change in protein concentration similar to that observed in therapeutic plasma exchange (TPE) in humans. The post-donation expression profile shows an increase in many proteins after even a single donation, and the effect is enhanced with additional donations. Unlike TPE, human plasma donation returns saline without albumin to the donor, thus eliminating supplemental albumin as a contributing factor. The observed fold change falls outside the variation expected by natural circadian expression changes.

N=6 humans (four donors, one monthly control and one circadian test). Seventeen antibody tests were performed, measuring the concentration of 507 proteins for each test.

## 1. Introduction

The interlinking of circulatory systems between young and old animals has been studied for more than a century. It has long been observed that the health of an older animal undergoing heterochronic parabiosis seems to improve after prolonged exposure to young blood. This effect has been the subject of intense research for the last few decades in the context of human longevity and healthspan, and the search for relevant blood factors is ongoing.

In May 2020, Mehdipour et al^1^. demonstrated that Neutral Blood Exchange (NBE), or the dilution of plasma using saline and albumin protein, was as effective as heterochronic parabiosis in altering circulating protein expression in mice. The post-NBE mice also showed improvements in muscle repair, liver adiposity and fibrosis, and a number of other key indicators of health. Similar changes in protein concentration were demonstrated in three human participants before and after therapeutic plasma exchange (TPE). While the dilution of plasma lowered the concentration of several proteins associated with an older phenotype one month after treatment, it also *increased* concentration of proteins associated with a younger phenotype. This sustained increase in concentration after dilution implies that a regulatory change was induced by TPE.

Therapeutic plasma exchange was chosen by the authors for practical reasons:

> *“*…*we took advantage of the fact that there is a procedure for human patients analogous to NBE where most of the plasma is replaced by physiologic solution, supplemented with commercial human albumin, called Therapeutic Plasma Exchange, TPE, which is FDA approved and routinely used in the clinic*.*”*

This was followed up with additional work on NBE and TPE demonstrating improved cognition and attenuated neuroinflammation in mice, as well as further analysis of human proteomics related to brain function pre- and post-TPE^2^. This surprising and consistent effect further led the authors to suggest that TPE should be considered as a rejuvenation therapy for removing deleterious age-related blood factors^3^.

### 1.1 TPE vs. commercial plasmapheresis

First successfully used therapeutically in 1952^4^, TPE has been shown to be effective in a variety of important clinical applications, from the removal of pathogenic circulatory factors such as antibodies, cytokines, triglycerides^3^, and endotoxins^4^. Blood is removed from the patient and plasma is separated by centrifuge. Red blood cells are returned to the body along with a solution of saline, albumin, and an anticoagulant factor such as sodium citrate. An amount typically equivalent to 1-1.4 times the patient’s estimated plasma volume (EPV) is exchanged for each treatment. The total number of treatments depends on the condition being treated, the concentration, molecular weight, and rate of regeneration of the factors to be removed^5^.

Commercial plasma donation (plasmapheresis) functions in a similar manner, with a few important differences. The plasma is exchanged with saline and anticoagulant without the addition of albumin, and the total amount of plasma removed from the body during plasmapheresis is relatively small (0.15-0.2 * EPV). Several plasma donations would be required to achieve the same volume of plasma exchange as TPE.

### 1.2 Approximating TPE with plasma donation

In Mehdipour et al, the mice treated with NBE had approximately half of their plasma exchanged, while a significant shift in protein profile was observed in humans after a single TPE treatment. While the total plasma volume removed by TPE was not disclosed, it would typically be much higher than 0.5 EPV.

Until the specific mechanism behind this effect is understood and the relevant blood factors are identified, it is not possible to estimate a rate of recovery or calculate an optimal donation schedule to dilute or remove them. Ignoring recovery, we can estimate the number of donations required to achieve 50% total dilution. This is illustrated in **Figure 1**.

**Figure 1:**
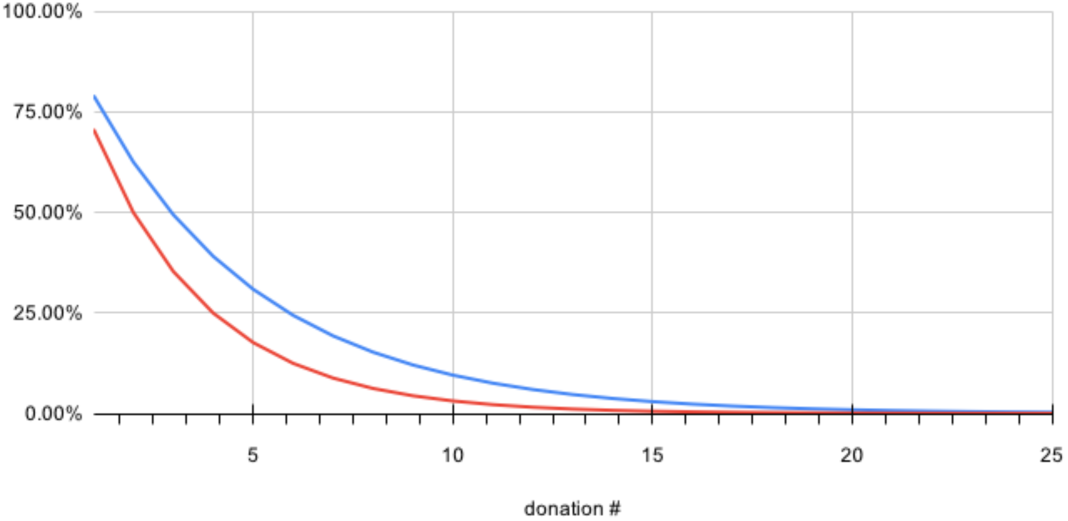
Estimated plasma dilution rates for 3000 mL (blue) and 4200 mL (red) EPV at 880 mL per donation

While the FDA permits plasma donation up to twice per week^6^, commercial plasma donation centers may set their own more stringent limits. This is an important constraint on the total rate of plasma that may be exchanged by a healthy individual in the United States.

If 50% dilution is sufficient to see change in expression, this can be achieved in most people with only 2 to 3 plasma donations.

## 2. Our experiment

We isolated plasma from EDTA-treated blood acquired via diabetic lancet from six individuals, and measured the concentration of proteins using an antibody detection array.

Four people (participants **A-D**) donated plasma at a commercial plasma donation center over a period of three months. Donations were made at various intervals ranging from weekly to bimonthly. The amount of plasma removed for commercial donation is determined by the provider following FDA guidelines, and typically represents 0.2-0.3 EPV. Plasma protein concentrations were tested roughly five days after donation.

One person (participant **E**) served as a control over the same three month period, providing blood test samples without plasma donation.

One person (participant **F**) provided blood test samples three times in a single day to measure the effect of circadian protein expression variation. Tests were performed in the early morning, afternoon, and late evening with roughly 7 hours between tests. This participant did not donate plasma.

The specific demographics of each participant are provided in **Table 1**. The cohort age range was 34-46.

**Table 1:**
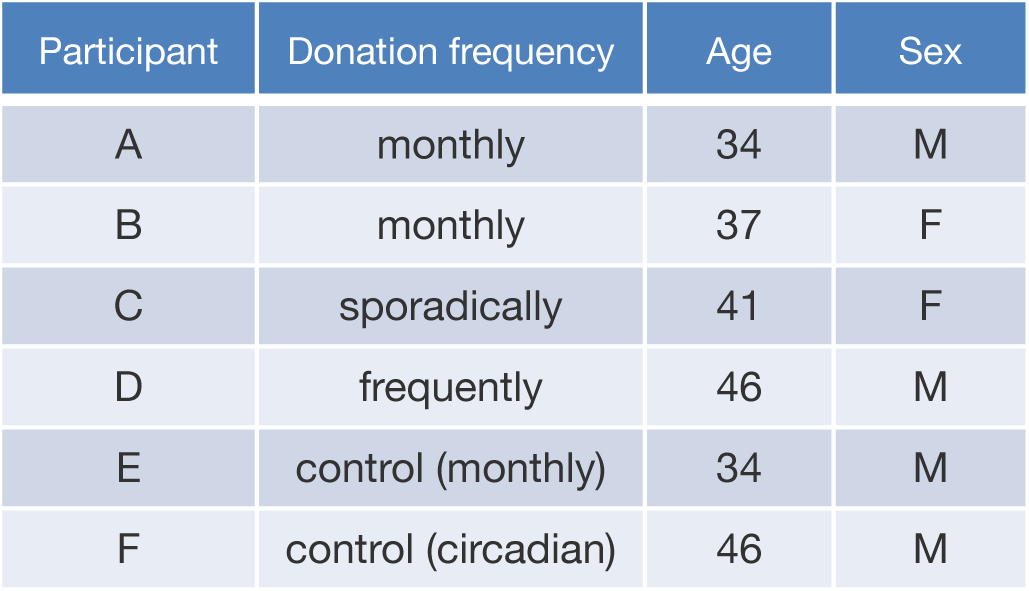
Participant demographics

| Participant | Donation frequency | Age | Sex |
| --- | --- | --- | --- |
| A | monthly | 34 | M |
| B | monthly | 37 | F |
| C | sporadically | 41 | F |
| D | frequently | 46 | M |
| E | control (monthly) | 34 | M |
| F | control (circadian) | 46 | M |

### 2.1 Post-donation results

All donors showed a clear *increase* in concentration of several proteins after plasma donation. The effect was more profound after additional donations. **Figure 2** shows a heatmap of relative expression of the top 30 proteins with significant change in at least one donor after donation.

**Figure 2:**
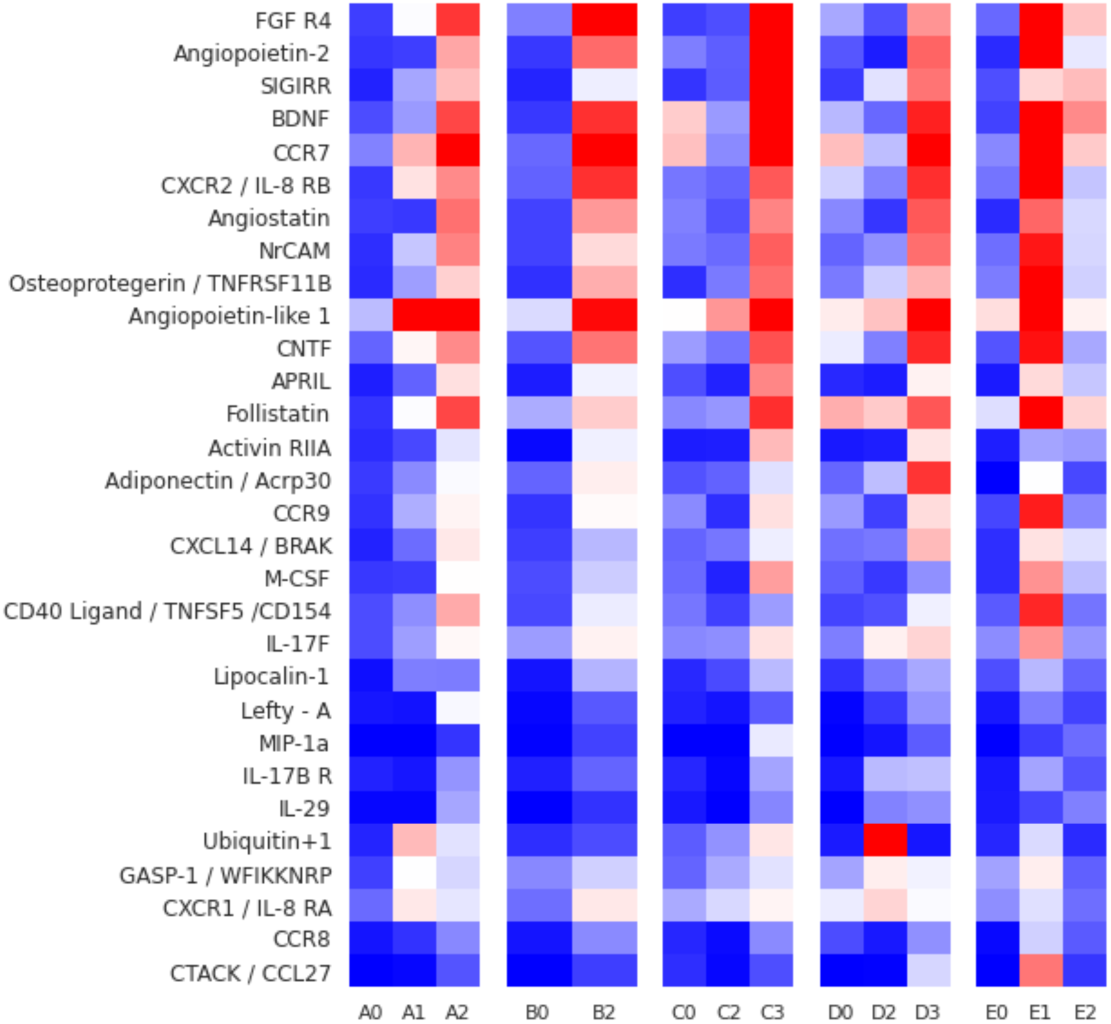
The top 30 proteins showing an increase in concentration after donation vs. the control.

Relative protein concentrations are shown, including intermediate donations. Each column represents a different participant, and cells within columns are colored blue-to-red (lowest-to-highest) per participant.

The first four columns show plasma donors. The first cell in each donor column shows the baseline concentration prior to donation. Rows that change from blue to red across each column indicate an increase concentration over time.

The final column shows samples from control participant **E** taken over the same period. The second cell in the control column shows the highest concentration of many proteins, with no clear trend across rows.

The donation schedule for each participant is listed in **Table 2** in the Supplementary Materials. Participants A and B followed a monthly schedule, with more frequent donations for C and D.

**Table 2:**
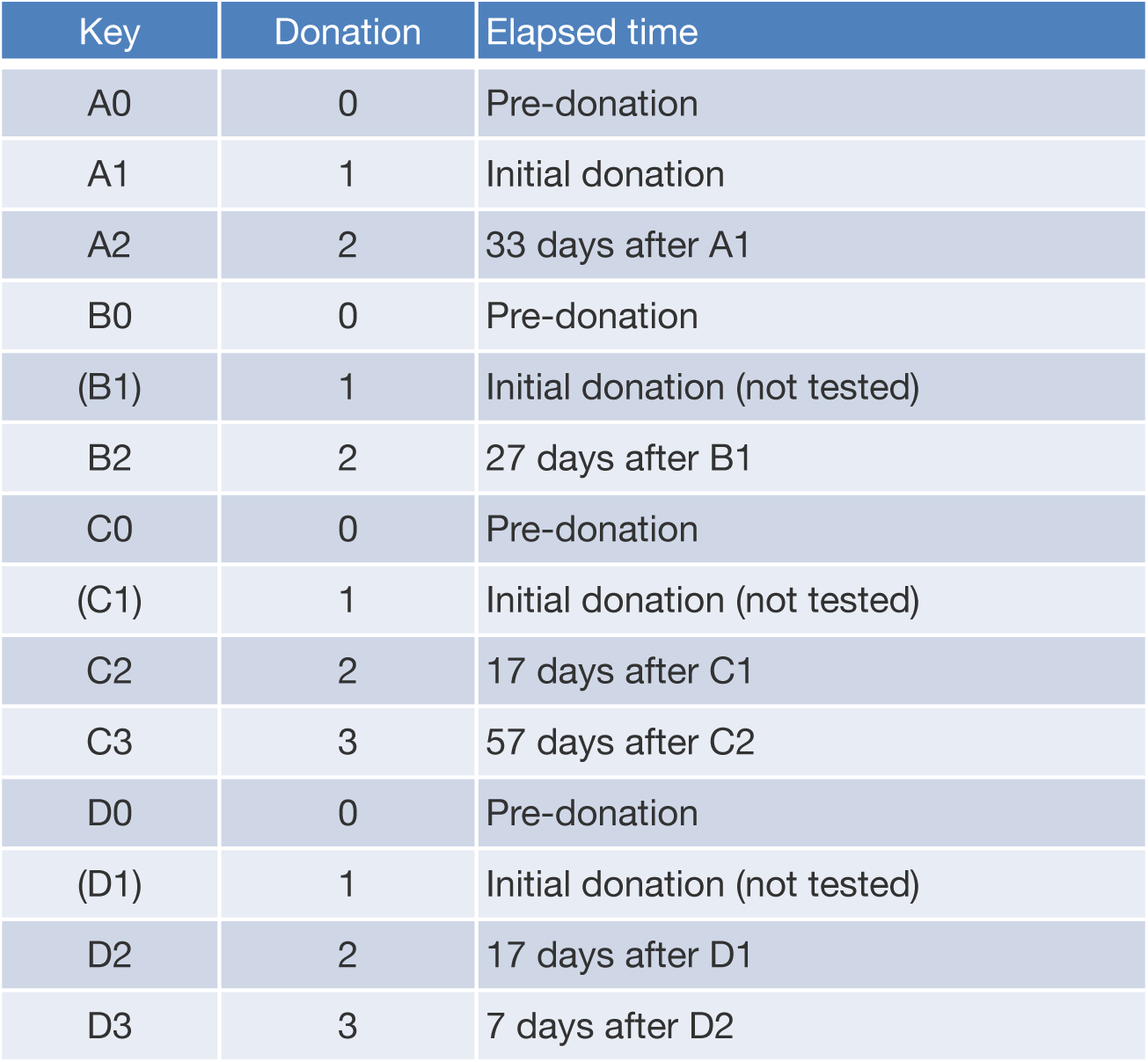
Participant donation schedule

The change in expression after TPE shown in Figure 5E of Mehdipour et al^1^. is recreated in **Figure 3** using data from the four plasma donors. Samples are shown in all before / all after order for the original 53 proteins of interest. While most proteins show a change after plasma donation similar to that observed after TPE in the original study, the effect is not consistent across all proteins or all samples in our cohort. A few (such as **Glut2**) even show a negative correlation.

**Figure 3:**
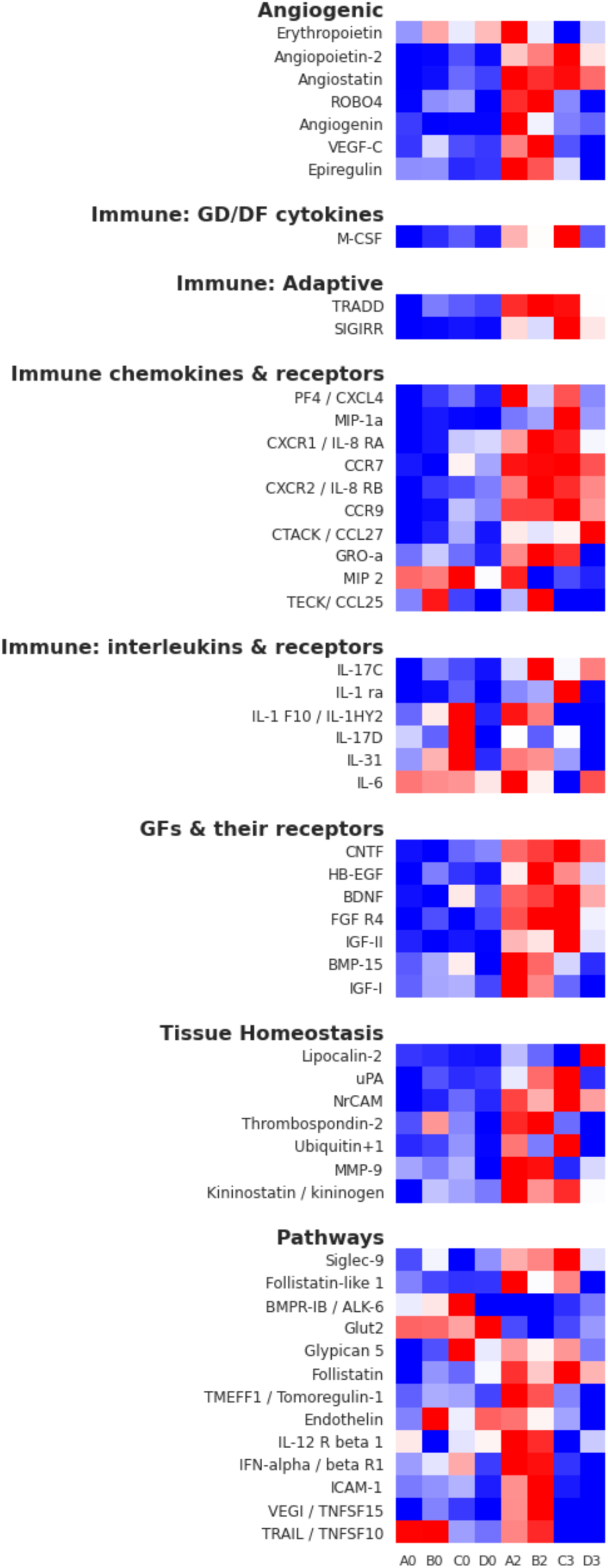
Pre- and post-donation concentrations in all before / all after order.

### 2.2 Change due to circadian variation

Human protein expression varies throughout the day^7^, and these daily expression profiles evolve throughout an individual’s lifespan^8^. The natural fluctuation of protein concentrations should be considered when measuring comparative protein levels.

Participant **F** provided blood test samples three times in a single day to measure the effect of circadian protein expression variation. Tests were performed in the early morning, afternoon, and late evening with approximately 7 hours between tests. This participant did not donate plasma. **Figure 4** (see Supplementary Materials) shows the fold increase or decrease compared to the morning baseline for all proteins.

**Figure 4:**
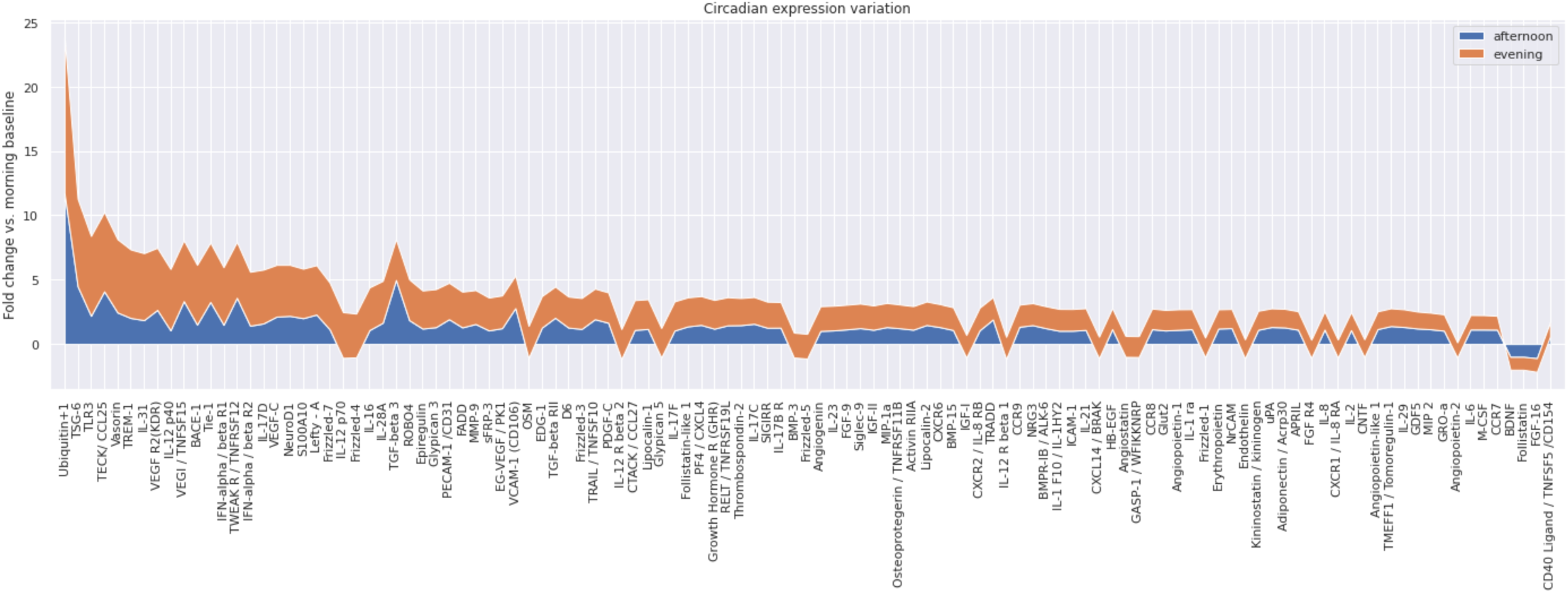
Fold increase or decrease of afternoon and evening samples vs. morning baseline in participant F.

All donor plasma tests were performed in the early afternoon. While the time awake prior to testing was not carefully controlled for every participant, the median rate of change from afternoon to evening in the circadian experiment was approximately **0.27** fold per hour.

Observed fold change for afternoon and evening samples vs. the calculated rate change / hour are shown in **Figure 5**.

**Figure 5:**
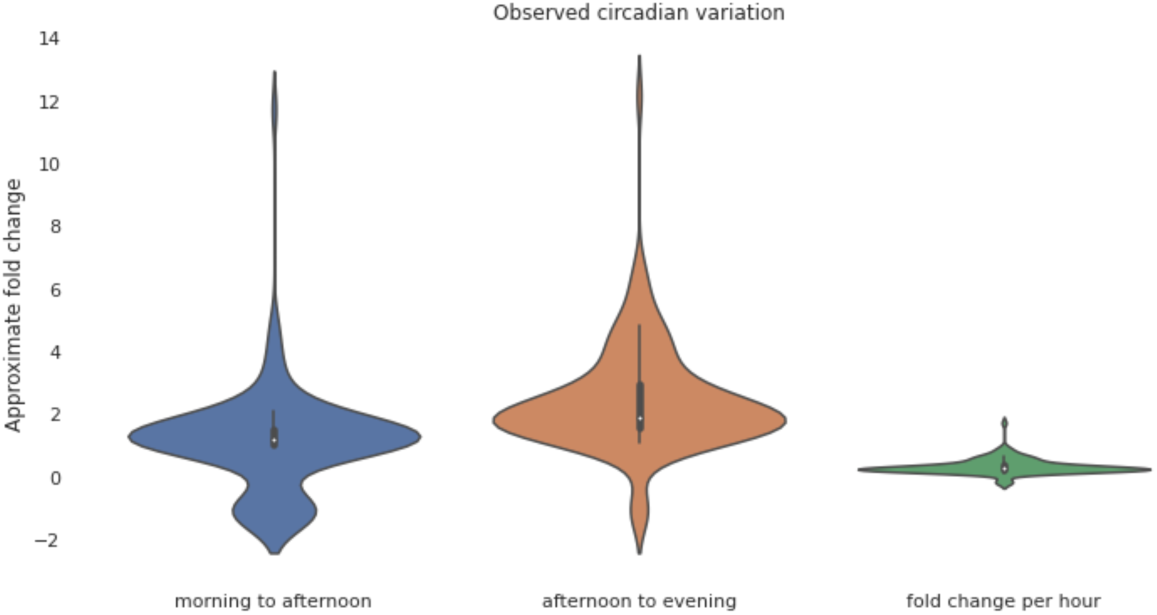
Observed fold change due to circadian variation.

Even allowing for a few hours variance in time awake between donors, this is well below the observed fold change for the top proteins of interest.

### 2.3 Top gains for all donors

While some proteins only show relative gains after more than one donation, twenty proteins show a consistent post-donation gain across all four donors after every donation. They are shown sorted by the median fold gain in **Figure 6**.

**Figure 6:**
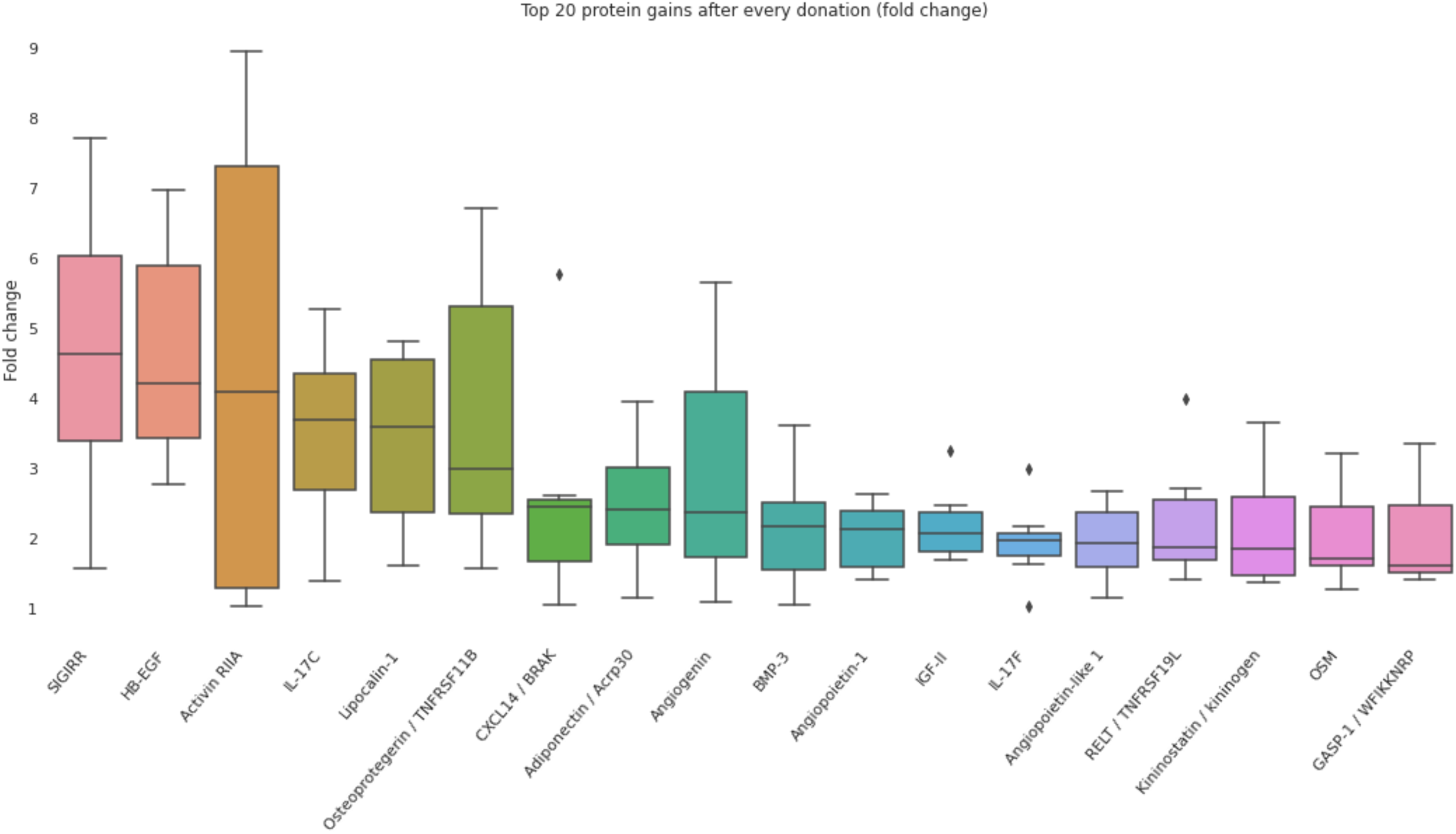
The twenty proteins showing a consistent post-donation fold increase across all donors.

## 3. Discussion

Many proteins show an increase in post-donation expression consistent with that observed after TPE, which has already been shown to be correlated with a younger phenotype. In addition, several previously unreported proteins also show a similar post-donation effect. A heatmap of these newly identified proteins are provided in **Figure 7** in the Supplemental Materials.

**Figure 7:**
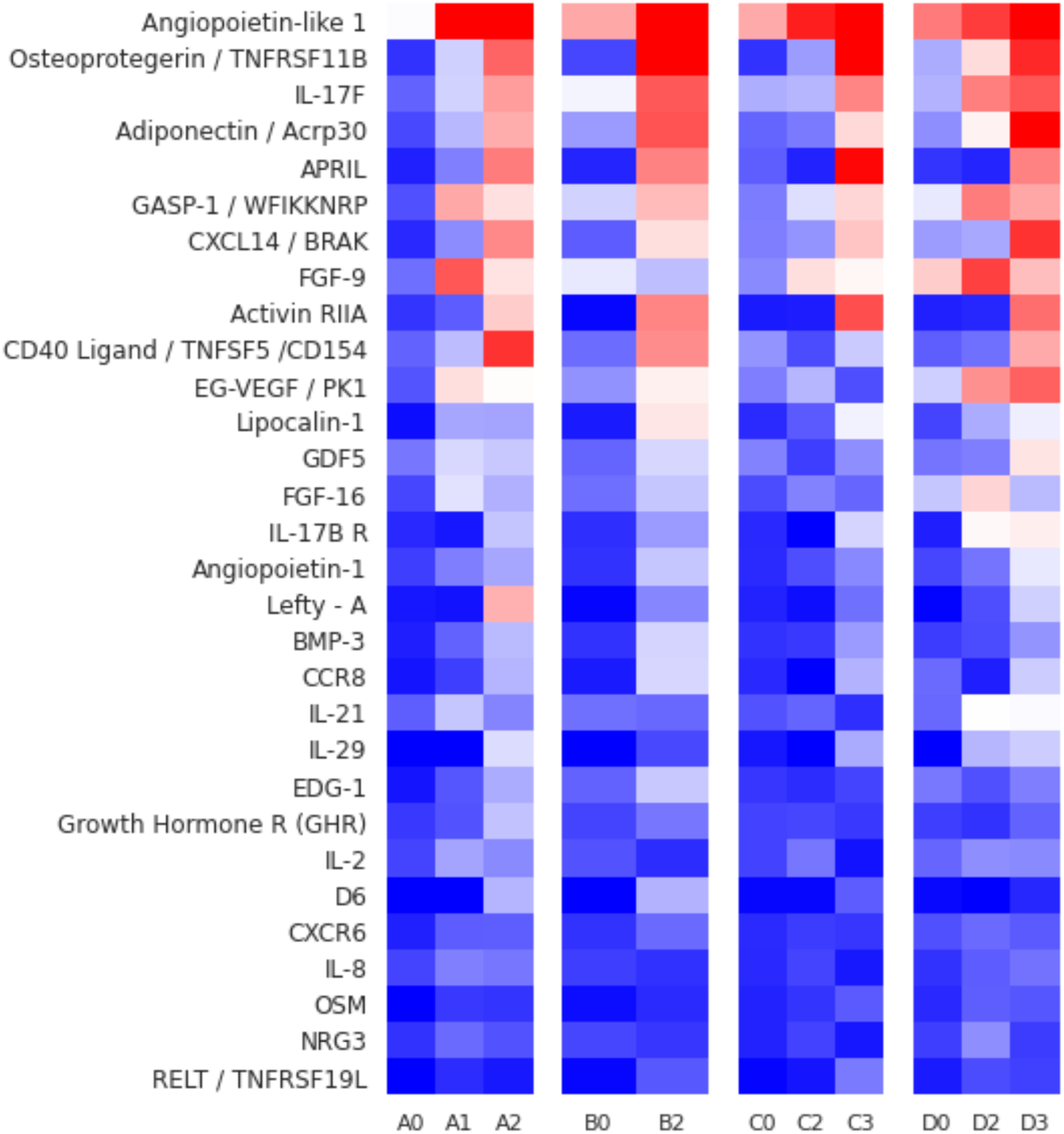
Additional proteins found to increase after plasma donation

While the expected change in plasma concentration due to natural circadian effects is relatively small in one hour, it can vary by 10 fold or more over the course of a day. While this circadian experiment was only performed for N=1 human, it would be prudent for any comparative proteomics test to control for awake time.

The underlying biochemical basis for this effect is currently the subject of intense study. The observed increase in concentration may be caused by the removal of inhibiting regulatory factors. But if the presence of such factors contributes to the march towards an older phenotype, why are they not naturally degraded? Commercial plasma donation is generally recognized as safe, and millions of liters are donated annually in the US. If the removal of these hypothetical down-regulators had a negative cumulative impact, it would surely have been observed by now in plasma donors. Finding such factors (or the underlying cause for their accumulation) and targeting them for intervention could be highly beneficial.

While our small commercial plasma experiment eliminates human albumin protein as a possible confounding factor, many questions remain. How long do these effects continue to be observed post-donation? If the duration of the effect proves to be significant, is this indicative of an epigenetic change? Does the anti-coagulant used (sodium citrate for plasmapheresis/TPE, heparin for mice undergoing NBE) play a significant role? Can the post-donation effect be shown in a larger cohort with more diversity in age and genetic background?

Commercial plasma donation is generally available to healthy adults in many countries, typically at no cost to the donor beyond the time required for donation. Some centers even pay the donor a modest fee for each donation. But despite the wide availability of commercial plasmapheresis, human plasma is a lifesaving resource consistently in short supply.

If this shifted expression profile continues to show rejuvenative potential in humans, it could provide motivation for eligible people to donate more frequently, and ultimately have a beneficial societal impact on donors and recipients alike.

## 4. Materials and methods

Protein tests were performed using a RayBiotech Human L507 Array^9.^. For each test, approximately 50uL of blood was extracted from a finger using a diabetic lancet and pipette, then transferred to a microcentrifuge tube containing 10uL EDTA. After centrifuging, plasma was extracted and prepared according to the manufacturer’s instructions. Plasma samples were frozen at -20C prior to preparation. Laser scanning and primary analysis of each prepared slide was performed by RayBiotech. Brightness data was normalized using the RayBiotech analysis tool.

Plots were made with Jupyter Notebook and Seaborn.

All code and data are available via GitHub: https://github.com/InterimResearch/plasma_donation

## Acknowledgements

The author would like to thank Adam Cecchetti, Mikhail Davidov, Lillian Johnson, Beth Kolko, Baron Oldenburg, Rich Olson, Isabella Pestovski, and Phebe Rossi for their contributions and feedback.

## Ethics declaration

The author declares that he has no competing interest. All participants were fully informed of the parameters of the experiment and provided written consent for publication. Each plasma donor performed donations at a commercial plasma donation facility following a self-chosen schedule.

## Supplementary materials

## References

1. Mehdipour, Melod, Colin Skinner, Nathan Wong, Michael Lieb, Chao Liu, Jessy Etienne, Cameron Kato, Dobri Kiprov, Michael J. Conboy, and Irina M. Conboy. “Rejuvenation of Three Germ Layers Tissues by Exchanging Old Blood Plasma with Saline-Albumin.” Aging 12, no. 10 (2020): 8790–8819. https://doi.org/10.18632/aging.103418.

2. Mehdipour, Melod, Taha Mehdipour, Colin M. Skinner, Nathan Wong, Chao Liu, Chia-Chien Chen, Ok Hee Jeon, Yi Zuo, Michael J. Conboy, and Irina M. Conboy. “Plasma Dilution Improves Cognition and Attenuates Neuroinflammation in Old Mice.” GeroScience 43, no. 1 (2020): 1–18. https://doi.org/10.1007/s11357-020-00297-8.

3. Mehdipour, Melod, Jessy Etienne, Chao Liu, Taha Mehdipour, Cameron Kato, Michael Conboy, Irina Conboy, and Dobri D. Kiprov. “Attenuation of Age-Elevated Blood Factors by Repositioning Plasmapheresis: A Novel Perspective and Approach.” Transfusion and Apheresis Science 60, no. 3 (2021): 103162. https://doi.org/10.1016/j.transci.2021.103162.

4. Clark, W.F., and S.S. Huang. “Introduction to Therapeutic Plasma Exchange.” Transfusion and Apheresis Science 58, no. 3 (2019): 228–29. https://doi.org/10.1016/j.transci.2019.04.004.

5. Kaplan, Andre A. “Therapeutic Plasma Exchange: Core Curriculum 2008.” American Journal of Kidney Diseases 52, no. 6 (2008): 1180–96. https://doi.org/10.1053/j.ajkd.2008.02.360.

6. “Plasmapheresis.” Code of Federal Regulations, title 21 (2015): https://www.accessdata.fda.gov/scripts/cdrh/cfdocs/cfcfr/CFRSearch.cfm?CFRPart=640showFR=1subpartNode=21:7.0.1.1.7.7

7. Braun, Rosemary, William L. Kath, Marta Iwanaszko, Elzbieta Kula-Eversole, Sabra M. Abbott, Kathryn J. Reid, Phyllis C. Zee, and Ravi Allada. “Universal Method for Robust Detection of Circadian State from Gene Expression.” Proceedings of the National Academy of Sciences 115, no. 39 (2018). https://doi.org/10.1073/pnas.1800314115.

8. Lehallier, Benoit, David Gate, Nicholas Schaum, Tibor Nanasi, Song Eun Lee, Hanadie Yousef, Patricia Moran Losada, et al. “Undulating Changes in Human Plasma Proteome Profiles across the Lifespan.” Nature Medicine 25, no. 12 (2019): 1843–50. https://doi.org/10.1038/s41591-019-0673-2.

9. RayBiotech Human L507 Array, https://www.raybiotech.com/l-series-507-label-based-human-array-1-glass-slide-2/

